# Dynamic coupling of cochlear inner hair cell intrinsic Ca^2+^ action potentials to Ca^2+^ signaling of non-sensory cells

**DOI:** 10.1101/731851

**Authors:** Federico Ceriani, Stuart L. Johnson, Miloslav Sedlacek, Aenea Hendry, Bechara Kachar, Walter Marcotti, Fabio Mammano

## Abstract

The relationship between Ca^2+^ action potential (AP) activity in immature inner hair cells (IHCs) and the spontaneous ATP-dependent intercellular Ca^2+^ signaling in cochlear non-sensory cells (NSCs) of the greater epithelial ridge (GER) is unclear. Here, we determined that IHCs fired asynchronous Ca^2+^ APs also in the absence of Ca^2+^ activity in the GER. Patch clamp recordings from IHCs isolated from the rest of the sensory epithelium confirmed that this firing activity is an intrinsic property of immature IHCs. However, frequency, correlation index and burst duration of IHC APs increased significantly during Ca^2+^ wave propagation in NSCs, and depended on wave extension in the GER. Furthermore, IHC depolarization under whole cell patch clamp conditions triggered Ca^2+^ signals in nearby NSCs with a delay that was proportional to the distance from the stimulated IHC. Thus the immature mammalian cochlea supports bidirectional exchange of Ca^2+^ signals between IHCs and NSCs.

**IMPACT STATEMENT:** In inner hair cells of the developing mammalian cochlea, Ca^2+^ action potentials are both intrinsic and bidirectionally coupled to the ATP-dependent Ca^2+^ signaling of the surrounding non-sensory cells.

## INTRODUCTION

The primary sensory receptors of the mammalian cochlea, the inner hair cells (IHCs), fire Ca^2+^-dependent action potentials (APs) during pre-hearing stages of development (Kros et al., 1998; Beutner and Moser, 2001; Marcotti et al., 2003a; Johnson et al., 2011; Johnson et al., 2012; Johnson et al., 2017), which occurs at around postnatal P12 in mice (Mikaelian and Ruben, 1965; Shnerson and Pujol, 1981). During the same time window, periodic transient elevations of cytosolic free Ca^2+^ concentration occur spontaneously in the greater epithelial ridge (GER) and propagate as intercellular Ca^2+^ waves invading variable portions of the GER (Tritsch et al., 2007; Schutz et al., 2010; Rodriguez et al., 2012). The relationship between the AP activity in IHCs and Ca^2+^ signaling in the GER is still not fully understood (Mammano and Bortolozzi, 2018).

The GER comprises the tall columnar cells of Kölliker’s organ (Kikuchi and Hilding, 1965; Hinojosa, 1977) and all other cells of the sensory epithelium that are located on the modiolar side of the pillar cells, including the inner phalangeal cells, the border cells and the IHCs (Kelley, 2007). Connexin 26 (Cx26) and connexin 30 (Cx30) are the prevailing isoforms expressed in NSCs of the GER (Forge et al., 2003; Mammano, 2019) and their block prevents the generation of Ca^2+^ waves in the GER (Schutz et al., 2010; Xu et al., 2017; Mammano and Bortolozzi, 2018). Furthermore, intercellular Ca^2+^ wave propagation is hampered in Sox10-Cre;*Gjb2*^fl/fl^ mice (Anselmi et al., 2008), with targeted ablation of Cx26 (Crispino et al., 2011), as well as in Cx30 homozygous knockout– LacZ mice (*Gjb6*^−/−^) (Anselmi et al., 2008; Ortolano et al., 2008; Rodriguez et al., 2012), both of which fail to acquire hearing (Teubner et al., 2003; Crispino et al., 2011; Fetoni et al., 2018). In contrast, intercellular Ca^2+^ waves propagate normally in mice that are global knockouts for pannexin 1 (*Panx1*^*–/–*^) (Zorzi et al., 2017; Mammano, 2019), and have hearing thresholds indistinguishable from those of littermate controls (Abitbol et al., 2019). Altogether, this evidence indicates that Ca^2+^ waves in the GER rely on ATP release from connexin hemichannels in the apical surface of NSCs (Tritsch et al., 2007; Schutz et al., 2010; Rodriguez et al., 2012; Xu et al., 2017; Mammano and Bortolozzi, 2018; Ceriani et al., 2019).

The ATP released during Ca^2+^ wave propagation has been proposed to trigger AP activity in immature IHCs, driving bursts of APs in the auditory nerve fibres (Tritsch et al., 2007). Extracellularly applied ATP caused elevation of cytosolic Ca^2+^ in adult IHCs (Dulon et al., 1991), due to activation of both ionotropic (P2X) and metabotropic (P2Y) purinergic receptors (Sugasawa et al., 1996). Immunolabeling experiments have demonstrated the presence of P2X_2_ purinergic receptors in the apical surface of IHCs, and shown to promote cell depolarization when activated by high concentrations of extracellular ATP (mature IHCs: Sugasawa et al., 1996; immature IHCs: Johnson et al., 2011). In contrast, ATP concentrations in the submicromolar range caused hyperpolarization in immature and mature IHCs, and this effect has been ascribed to a functional coupling between P2X receptors and SK2 channels (Sugasawa et al., 1996; Johnson et al., 2011).

Here, we used confocal Ca^2+^ imaging and patch clamp electrophysiology to provide an in depth investigation of the relationship between spontaneous AP activity in IHCs and intercellular Ca^2+^ waves in NSCs of the GER. Our results indicate that ATP-dependent signaling in cochlear NSCs is not required for AP generation in pre-hearing IHCs, but can increase the frequency of Ca^2+^ spikes in IHCs, thus increasing the probability of synchronized firing among nearby sensory cells.

## MATERIALS AND METHODS

### Ethics Statement

In the UK, experiments were performed in accordance with Home Office regulations under the Animals (Scientific Procedures Act) 1986 and following approval by the University of Sheffield Ethical Review Committee. In Italy, all experiments involving the use of animals (mice) were performed in accordance with a protocol approved by the Italian Ministry of Health (Prot. n.1276, date 19/01/2016). In the USA, experiments were conducted in accordance with the Guide for the Care and Use of Laboratory Animals by NIH and approved the Animal Care and use Committees for the National Institute on Deafness and Other Communication Disorders (NIDCD ACUC, protocol #1215).

### Generation and genotyping of transgenic mice

For calcium imaging experiments in which genetically encoded calcium indicators were used, transgenic mice expressing GCaMP3 or GCaMP6f on the C57BL/6 background were utilized, bred to homozygosity and crossed with animals heterozygous for Gfi1-Cre (Yang et al., 2010) (hair cell only expression; MGI:4430258) or Emx2-Cre (Kimura et al., 2005) (whole epithelium expression; MGI:3579416). GCaMP3 knock-in mice (R26-lsl-GCaMP3) were a generous gift from Dwight E. Bergles (Johns Hopkins University), GCaMP6f mice (Madisen et al., 2015) were obtained from The Jackson Laboratory (B6;129S-*Gt(ROSA)26Sor*^*tm95*.*1(CAG-GCaMP6f)Hze*^/J; Stock No.:024105; MGI:5558088). Transgenic animals were genotyped by standard PCR using specific primers for GCaMP3 and GCaMP6f animals, and generic Cre primers to genotype for Gfi and Emx2.

### Tissue preparation

Ca^2+^ signaling and IHC activity in apical coil IHCs from wild type C57BL6/J mice of either sex were studied in either acutely dissected cochleae or in organotypic cultures from postnatal day 4-5 (P4-P5) pups, where the day of birth is P0. For organotypic cultures, cochleae were dissected in ice-cold Hepes buffered (10 mM, pH 7.2) HBSS, placed onto glass coverslips coated with 185 μg/ml of Cell Tak (Becton Dickinson) and incubated overnight at 37°C in DMEM/F12 (Thermo Fisher Scientific) supplemented with FBS 5%. Acute preparations were dissected in normal extracellular solution contained (in mM): 135 NaCl, 5.8 KCl, 1.3 CaCl_2_, 0.9 MgCl_2_, 0.7 NaH_2_PO_4_, 5.6 D-glucose, 10 Hepes-NaOH, 2 sodium pyruvate, amino acids and vitamins (pH 7.5; osmolality ∼308 mmol kg^-1^). Dissected cochleae were transferred to a microscope chamber, continuously perfused with a peristaltic pump using normal extracellular solution (see above) and viewed using an upright microscope (for microscope details, see Ca^2+^ imaging, below). Crucial for the generation of the AP activity in IHCs is the use of near physiological recordings conditions, which for the mammalian IHCs includes working at body temperature (35-37°C) and using a perilymph-like extracellular solution (1.3 mM Ca^2+^ and 5.8 mM K^+^) (Johnson et al., 2008; Johnson et al., 2017).

For experiments requiring the use of transgenic animals expressing Ca^2+^ biosensors, cochleae were dissected at room temperature in DMEM/F12 (Thermo Fisher Scientific) containing 10% fetal bovine serum (FBS) and ampicillin (5 µg/ml). After dissection, the tissue was mounted on a 10 mm glass cover slip coated with collagen and maintained in culture in a glass bottom Petri dish with DMEM/F12 at 37°C overnight. If cultures were kept in the incubator for another day, culturing media was exchanged 24 hours post dissection.

### Electrophysiology

Patch pipettes were made from Soda glass capillaries (Harvard Apparatus Ltd, UK) and coated with surf wax (Mr Zoggs SexWax, USA) to minimize the fast transient due to the patch pipette capacitance. Recordings were made as previously described (Johnson et al., 2008; Corns et al., 2014; Johnson et al., 2017) using an Axopatch 200B (Molecular Devices, USA) or an Optopatch (Cairn Research, UK) amplifier. Data acquisition was controlled by pClamp software using Digidata boards (Molecular Devices, CA, USA). The cell-attached AP recordings were filtered at 10 kHz, sampled at 20 kHz and some were filtered offline at 500 Hz (8-pole Bessel) using Clampfit (Molecular Devices). Data analysis was performed using Origin software (OriginLab, Northampton, MA, USA). The pipette solution used for cell-attached recordings contained (in mM): 140 NaCl, 5.8 KCl, 1.3 CaCl_2_, 0.9 MgCl_2_, 0.7 NaH_2_PO_4_, 5.6 D-glucose, 10 Hepes-NaOH (pH 7.5). Hair cells spike occurrence was collected in peri-stimulus-time histograms. The Mini Analysis Program (Synaptosoft Inc. NJ, USA) was used to detect spike events in cell-attached recordings. The AP frequency was computed as the reciprocal of the mean interspike intervals (ISI) for each cell. To derive a continuous function for the firing rate vs. time, spike trains were smoothed by convolution with a Gaussian kernel (S.D. = 6 s).

### Ca^2+^ imaging

For calcium dye loading, cochlear cultures and acutely dissected preparations from wild type animals were incubated for 40 min at 37°C in DMEM/F12, supplemented with fluo-4 acetoxymethil (AM) ester (final concentration 16 μM; Cat. No. F14217, Thermo Fisher Scientific). The incubation medium contained also pluronic F–127 (0.1%, w/v), and sulfinpyrazone (250 μM) (Sigma, UK) to prevent dye sequestration and secretion (Di Virgilio et al., 1989; Bootman et al., 2013). Preparations were then transferred on the microscope stage and perfused with extracellular solution for 20 minutes before imaging to allow for de-esterification. Fluo-4 signals were recorded using either a custom built spinning disk confocal microscope (Ceriani et al., 2016b) or a commercial two-photon confocal microscope (Ceriani et al., 2019).

For spinning disk confocal microscopy, Fluo-4 fluorescence was excited by light emitted from a 488 nm diode laser (COMPACT-150G-488-SM; World Star Tech), filtered through a narrow band filter (FF02-482/18-25, Semrock/IDEX Health & Science, LLC, Rocherster, N.Y.), and projected onto the sample by reflection off a 45° dichromatic mirror (Di02-R488-25×36, Semrock/IDEX Health & Science). Fluorescence emission was filtered through a OD6 bandpass interference filter (535/43M, Edmund Optics). Confocal fluorescence images were formed by a water immersion objective (40×, NA 0.8, LUMPLAFL, Olympus, Tokyo, Japan) and projected on a scientific–grade CMOS camera (Edge, PCO AG, Kelheim, Germany) controlled by software developed in the laboratory. Image sequences were acquired continuously at 10 Hz frame rate with 100 ms exposure time. To synchronize image acquisition and patch clamp recordings, we sampled the 5 V pulse (FVAL) that signals active exposure of the CMOS camera (Canepari and Mammano, 1999).

For multiphoton Ca^2+^ imaging with Fluo-4, a laser-scanning confocal microscope (Bergamo II System B232, Thorlabs Inc., Newton, New Jersey, USA) equipped with a water immersion objective, (60×, 1.1 NA, LUMFLN60XW, Olympus, Japan) was coupled to a mode-locked laser system (Chameleon Ultra II, Coherent Inc., Santa Clara, California, USA) operating at *λ* = 920 nm, 80 MHz pulse repeat, 140 fs pulse width. Fluorescence emission was filtered through a bandbass filter (FF02-525/40-25, Semrock/IDEX Health & Science) and detected by a cooled GaAsp PMT (H7422-40, Hamamatsu) to form raster-scan images (768×768 pixels) at 20 Hz frame rate.

For fluorescence imaging of GCaMP Ca^2+^ biosensors, we used an upright Nikon microscope equipped with Nikon water immersion objectives (either a 16× 0.8 NA CFI75 LWD, or a 25×C 1.1 NA CFI75 Apochromat), a Yokogawa CSU-21 spinning disc head, and an Andor DU-897 camera. For higher numerical aperture imaging, we flipped upside down the glass coverslip with the attached culture inside the glass bottom Petri dish and used drops of Vaseline as spacers. The upside down cultures (apical surface towards the objective) were viewed in a Nikon TiE inverted fluorescence microscope equipped with a Nikon objectives (either a 60×C 1.27 NA CFI SR Plan Apochromat IR water immersion objective, or a 100× 1.45 NA CFI Apochromat Lambda oil immersion objective), a Yokogawa CSU-21 spinning disc head, and an Andor DU-897 camera. NIKON Elements software was used for image acquisition. For pharmacology experiments, cultures were incubated with respective reagents for at least 20 minutes before imaging and then imaged in continuous presence of the drugs. Reagents were prepared fresh from stock solutions before each experiment.

### Mathematical methods

Images were analyzed off-line using software routines written in MATLAB (Matlab R2011a, The Mathworks Inc.), Python (Python 2.7, Python Software Foundation) and ImageJ (NIH). Ca^2+^ fluctuations were quantified either as relative changes of fluorescence emission intensity

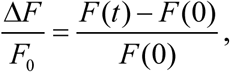

or as normalized increments

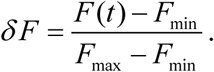

For the analysis in **Figure 3**, Ca^2+^ wave directions were estimated using an optical flow analysis algorithm (Afrashteh et al., 2017). First, each recording was preprocessed by applying a Gaussian (σ=2 pixels) and a median (2×2 pixels) filter. A Δ*F/F*_0_ image sequence was then generated from the raw data. Pixels belonging to different Ca^2+^ events were selected by drawing a ROI around the maximal extension of the wave. The processed recordings were then analyzed using Matlab and the Combined Local-global method of the Optical flow analysis toolbox (Afrashteh et al., 2017). We computed the distribution of instantaneous angles taken across all image frames of pixels within the ROIs. Finally, each wave was assigned to one of the three direction groups (IHCs to GER, GER to IHCs or longitudinal) based on the prevalent direction in the instantaneous angle distribution (IHC to GER: 225<α<315; GER to IHC: 45< α<135, longitudinal 135< α<225 and −45< α<45, where α is the angle between the optical flow vector and a line parallel to the IHCs, see **Figure 3A**).

**Figure 1.**
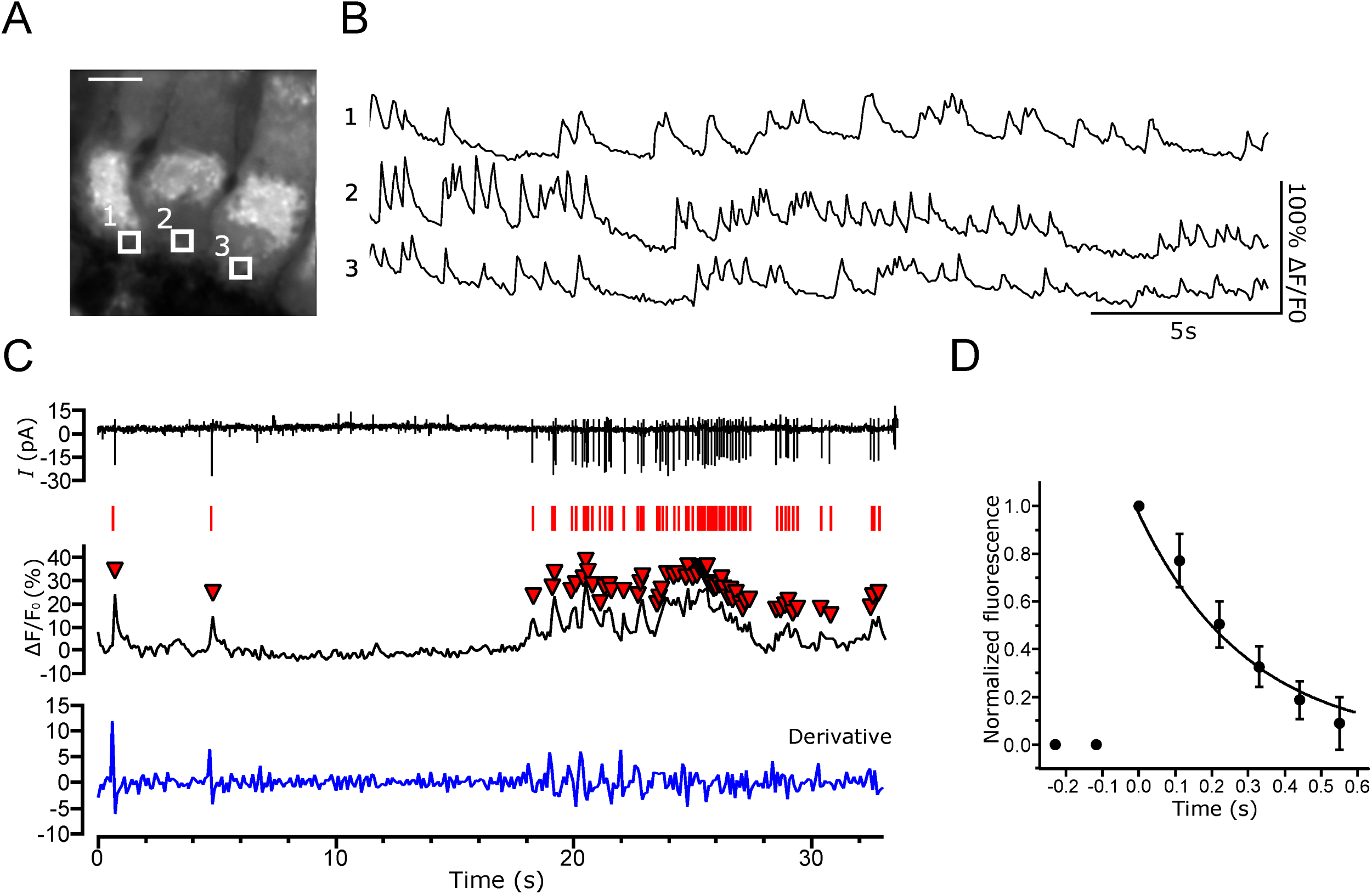
Optical detection of Ca^2+^-dependent action potentials in mouse cochlear IHCs. **(A)** High magnification image of P5 cochlear IHCs loaded with Fluo-4 AM (see also Supplementary Video 1). **(B)** Representative traces showing optical detection of Ca^2+^ signals from the synaptic region of three adjacent IHCs. Traces were generated as pixel average from the three ROIs highlighted in (A). **(C)** Comparisons of cell-attached patch clamp recordings from a P5 IHC (top) and fluorescence Ca^2+^ signals from the same cell (middle). The bottom trace represents the time derivative of the fluorescence signal. Red bars single out APs detected by high-pass filtering of the patch clamp trace; downward triangles denote (corresponding) peaks of the Δ*F*/*F*_0_ trace. **(D)** Estimate of decay time constant from normalized fluorescence signals associated to individual Ca^2+^ action potentials. Temperature 34-37°C.

**Figure 2.**
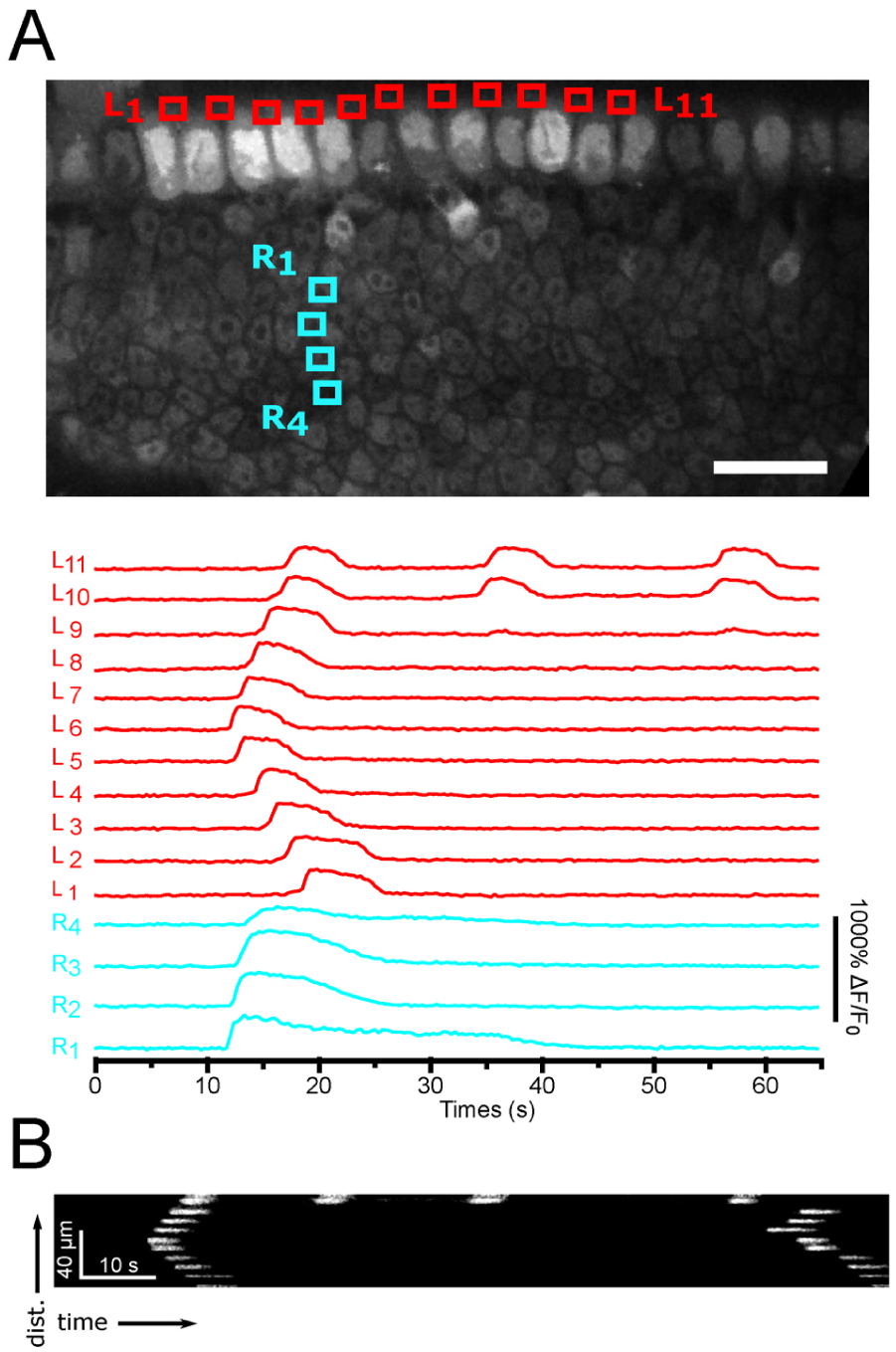
Ca^2+^ waves propagate across GER non-sensory cells. **(A)** Ca^2+^ waves that initiate in the proximity of inner phalangeal cells, which intercalate between IHCs, propagate both along the coiling axis of the cochlea (red ROIs and traces) and in the direction orthogonal to this (cyan ROIs and traces). **(B)** Kymograph corresponding to red traces in (A), showing repetitive Ca^2+^ waves that propagate along the L_1_ − L_11_ line. Note the nearly constant speed (slope) of these waves.

**Figure 3.**
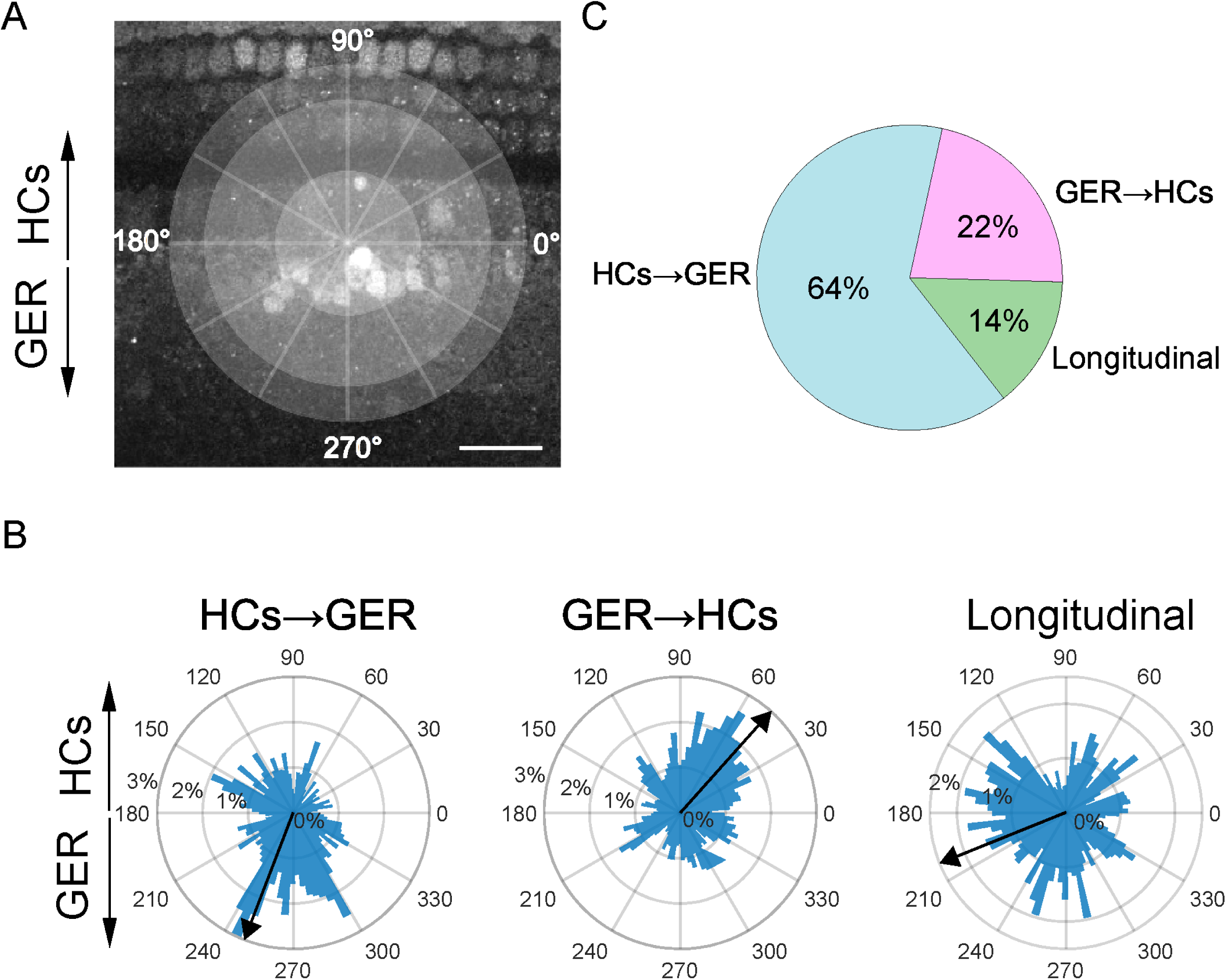
Directionality of Ca^2+^ waves in the GER. **(A)** Polar diagram superimposed to a reference fluorescence image of the organ of Corti. **(B)** Circle-chart showing the proportion of waves in each of the three directions (53 Ca^2+^ waves, 4 cochleae, 4 mice). **(C-E)** Polar histograms showing the distribution of instantaneous optical flow angles in three representative Ca^2+^ waves moving parallel to the tonotopic axis (**C**: longitudinal), from the IHC region towards the bulk of the GER **(D)** and from the GER towards the IHCs **(E)**.

For correlation analyses (**Figures 4-6**), Ca^2+^ waves were identified using thresholding, and a region of interest (ROI) was drawn around the maximum extension of each multicellular Ca^2+^ elevation event. Only events that remained confined to the field of view of the microscope were considered for this analysis.

**Figure 4.**
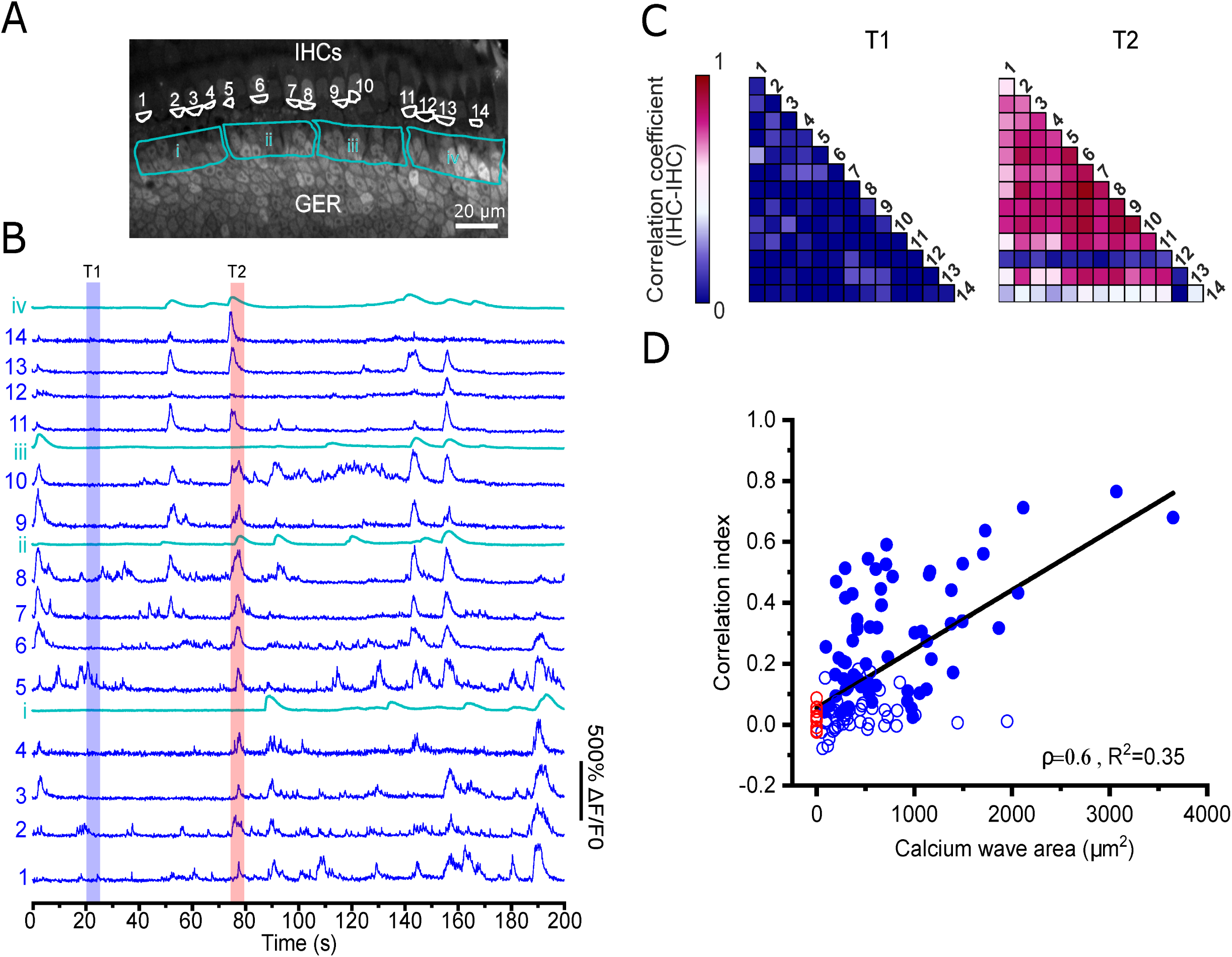
Relation between IHC Ca^2+^ action potential and Ca^2+^ waves in non-sensory cells in the GER. **(A**,**B)** Representative recording of Ca^2+^ signals from 14 IHCs (blue) and non-sensory cells from four adjacent zones in the GER (cyan). **(C)** Correlation matrices computed from the IHCs in panel ***B***. Correlation coefficients were computed in 5-s time windows in the absence (T1) or presence (T2) of Ca^2+^ waves in nearby non-sensory cells. **(D)** Correlation index between IHC activity as a function of Ca^2+^ wave area. Each data point represents the correlation index (average correlation coefficient) computed in a time window centered to the maximum amplitude of the Ca^2+^ wave (blue) or in the absence of Ca^2+^ waves in the GER (red). Filled dots: correlation index significantly different from 0 (P<0.05, Student’s t-test). Temperature 34-37°C.

**Figure 5.**
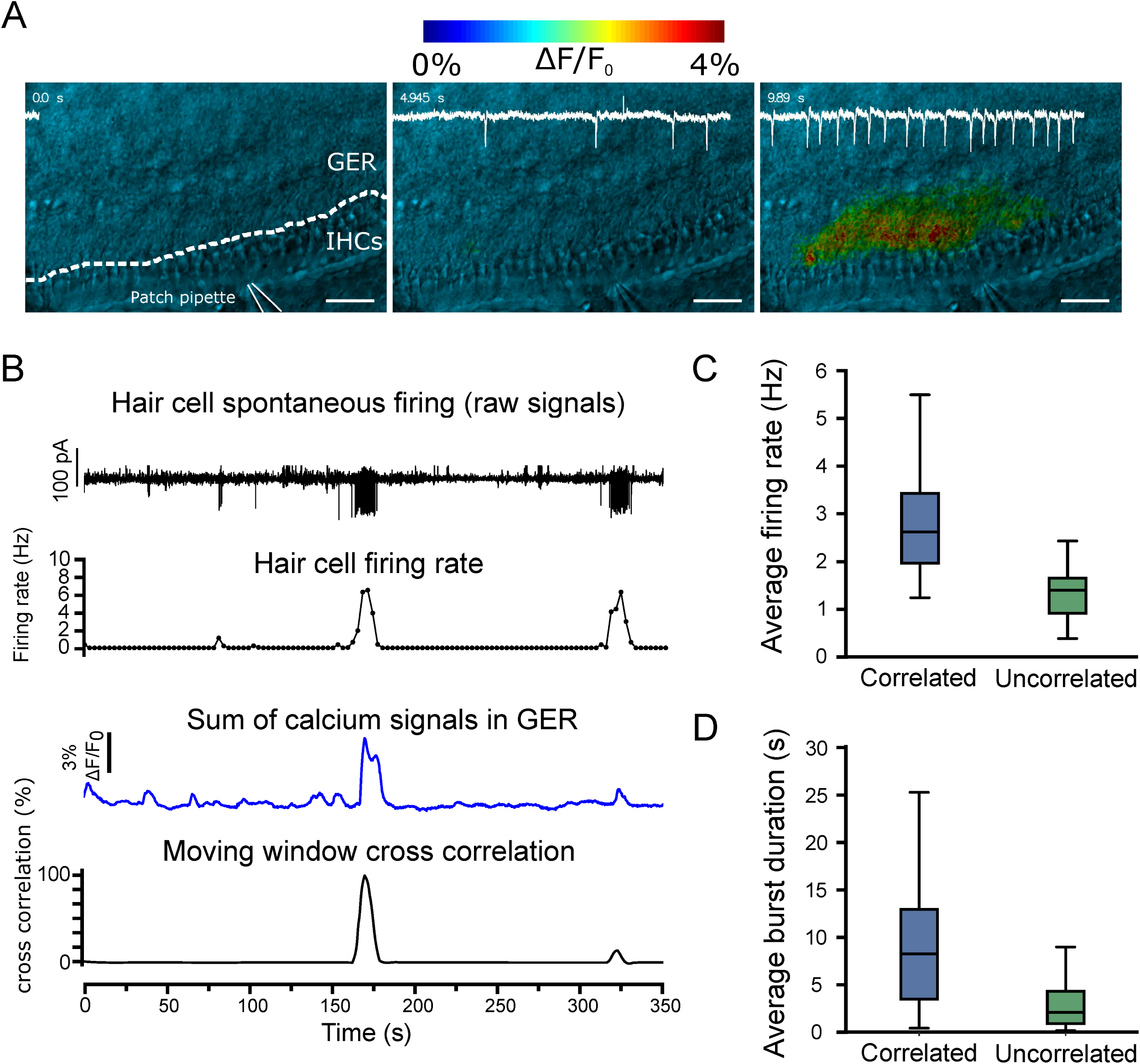
Modulation of IHC spontaneous activity by non-sensory cells. **(A)** Three representative frames from the same image sequence obtained while performing simultaneous Ca^2+^ imaging and cell-attached patch clamp recording from a P5 IHC. The top image shows the different regions of the sensory epithelium: the hair cell region (HCR) and the non-sensory cell region in the GER. The IHC was approached with the patch pipette from the outer hair cell side in order to avoid disruption the non-sensory cell in the GER. The IHC exhibited spontaneous electrical activity (white trace), the frequency of which increased when a Ca^2+^ wave was detected in nearby non-sensory cells (bottom panel). Scale bar: 25 μm. **(B-D)** Analysis of cross correlation between the IHC firing rate and Ca^2+^ signals in non-sensory cells. IHC burst frequency was significantly higher and burst duration was significantly longer when the firing was correlated with the occurrence of Ca^2+^ waves in proximal non-sensory cells.

**Figure 6.**
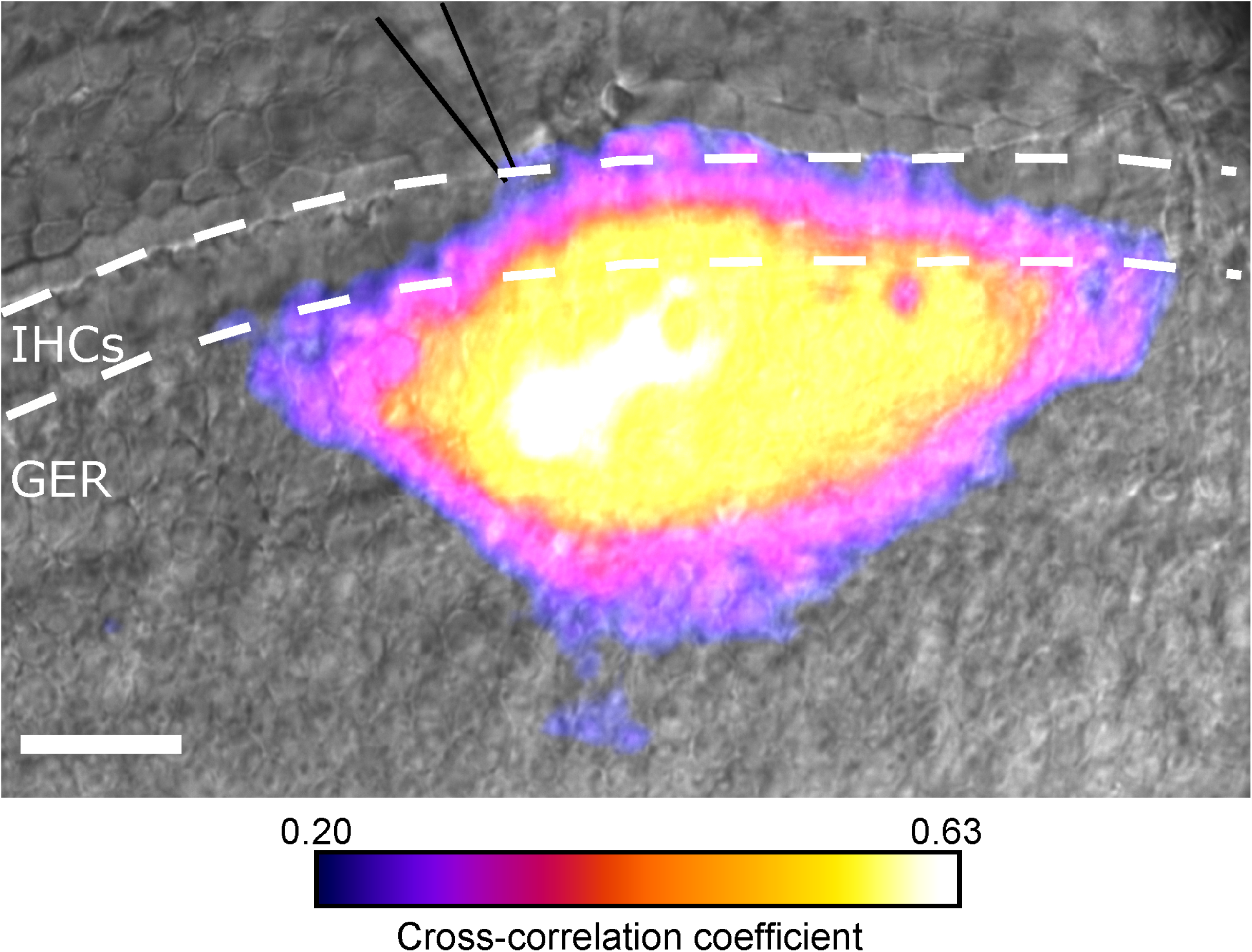
Spatial extent of IHC activity correlated with the occurrence of Ca^2+^ in the GER. Representative false-color image showing the spatial distribution of the cross correlation coefficient between Ca^2+^ transients in non-sensory cells and the firing rate of a patch-clamped IHC. Scale bar: 20 μm.

For the analysis in **Figure 4**, rolling (zero-lag) cross-correlation coefficients between different IHC Ca^2+^ traces were computed as:

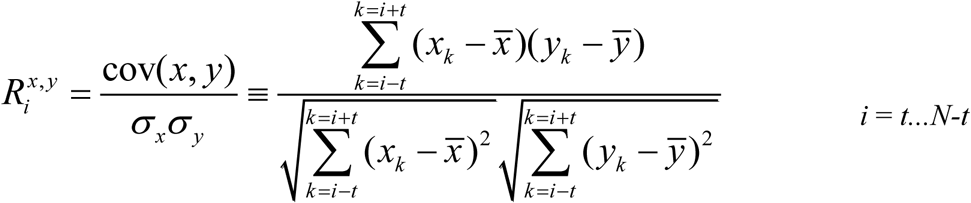

where *x* and *y* are two time series of size *N, 2t* is the size of the time window where the cross correlation was computed (5 seconds, or 100 frames) and 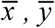 are the average values of *x* and *y* in the time window. To estimate the correlation index (*c*), we first computed the rolling cross correlation for every pair of identifiable IHCs in the field of view (on average 14±2 cells per recording) in a time window centered on the peak of an intercellular Ca^2+^ wave. We then averaged these values using Fisher’s *z*-transformation (Fisher, 1915): *z* = arctanh(*R_i_*), and its inverse *c* = tanh(*z*). Correlation index values are comprised between −1 and +1, with 0 indicating no correlation. This method was used also to estimate cross correlation coefficients between optical signals associated to GER Ca^2+^ waves and IHC firing rate, estimated from electrophysiological recordings (**Figures 5** and **6**). To this end, Ca^2+^ imaging datasets were first smoothed by applying a Gaussian (S.D. = 3 pixels) and median filter to every frame. Next, a baseline reference image (*F*_0_) was computed by averaging the first 5 frames of each recording. Time series were then generated by computing the pixel-by-pixel relative intensity change, Δ*F*/*F*_0_. To estimate the area of the GER were Ca^2+^ wave activity was significantly correlated to the spontaneous electrical activity of the patched IHC, we first down-sampled the IHC firing rate to the sample rate of the Ca^2+^ imaging acquisition. Then, a cross-correlation image was generated by calculating the cross correlation coefficient between the firing rate and the fluorescence values pixel-by-pixel. The background of this image was computed from an area of the GER that did not show any Ca^2+^ wave during the length of the recording. The longitudinal and radial extension of the correlation area was estimated using ImageJ.

For the analysis in **Figure 5**, IHC AP activity was considered significantly correlated to GER Ca^2+^ signaling activity when the cross-correlation coefficient was higher than a threshold that was determined using a Monte-Carlo approach by computing the (rolling) cross correlation of two pseudo-random traces having the same noise distribution as the original traces. The maximum value of the cross correlation coefficient was computed for 1000 different pairs of random traces, and the threshold was set at the 95% percentile.

For the analyses of GCaMP fluorescence signals in **Figures 8** and **9**, IHC activity was imaged and acquired using NIS-Elements software (Nikon Instruments, Inc.) at 0.5 – 2 Hz frame rate and analyzed later offline. Calcium transients were measured at the basal pole of contiguous IHCs, where single ROIs were manually set, as demonstrated in **Figure 8B**. Changes in fluorescence in IHCs during individual events were calculated as fluorescence of each event divided by the sum of total background fluorescence of the calcium indicator and event fluorescence (F_event_/F_total_). Fluorescence profiles of each ROI/IHC were then plotted over time in NIS-Elements and numbers extracted to Igor Pro (Wavemetrics), where the fluorescence signals were translated into traces of raw fluorescence as shown in **Figure 8D**. Next, calcium transients were detected using a custom written derivative thresholding routine in Igor Pro that marked each detected event (see also (Zhou et al., 2018). Accuracy of detected events was verified by visual inspection. Frequency of events was calculated for each individual IHC as total number of detected events during the duration of a single trial and converted to frequency in Hz. Amplitudes were further analyzed and plotted in distribution histograms (0.02 F_event_/F_total_ bins) for each experimental and pharmacological condition (**Figure 9A-C)** and normalized relative to control (**Figure 9D**).

**Figure 7.**
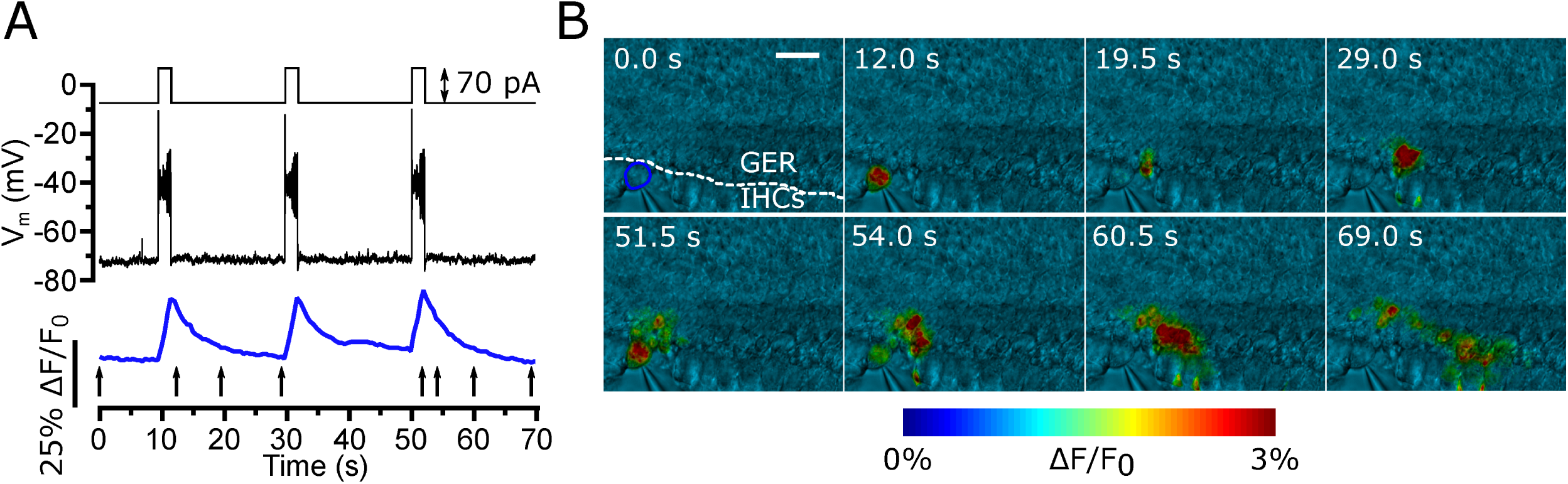
IHC depolarization triggers Ca^2+^ waves in GER non-sensory cells. **(A)** Representative recording of membrane potential (black trace) and Ca^2+^ signals (blue trace) from an IHC a from a P5 mouse cochlea triggered by a series of 70 pA current injections under current clamp (2 s in duration, inset: see **Materials and Methods**) from the IHC resting membrane potential. **(B)** Representative frames from the same recording in panel **A** showing Ca^2+^ waves in proximal cochlear non-sensory cells of the GER triggered by each IHC depolarization. The temporal position of each frame is indicated by black arrows in panel **A**. Experiments were performed at 20-22°C in order to prevent or minimize spontaneous action potentials activity in IHCs.

**Figure 8.**
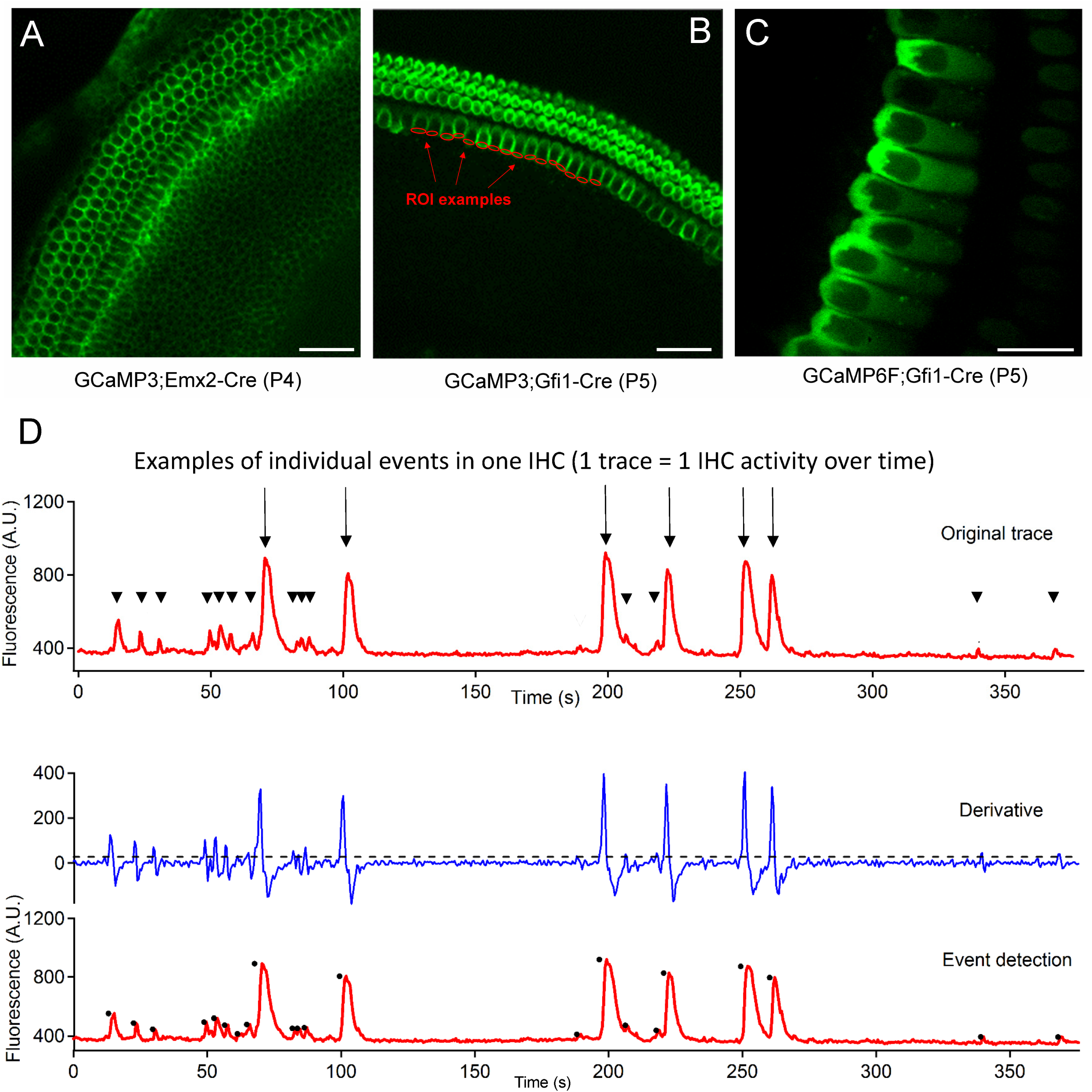
Expression and function of Ca^2+^ biosensors in the developing mouse cochlea. **(A-C)** Representative confocal fluorescence images of cochlear culture from GCaMP3;Emx2-Cre mice (**A**), GCaMP3;Gfi1-Cre mice (**B**) and GCaMP6f;Gfi1-Cre mice (**C**); scale bars: 40 µm in (**A)** and (**B)**, 40 µm in (**C**). **(D)** Detection of Ca^2+^ signals (events) from the basal pole of IHCs (see ROI examples in panel **B**) in one of these cultures based on time derivative computation of the raw fluorescence trace followed by thresholding.

**Figure 9.**
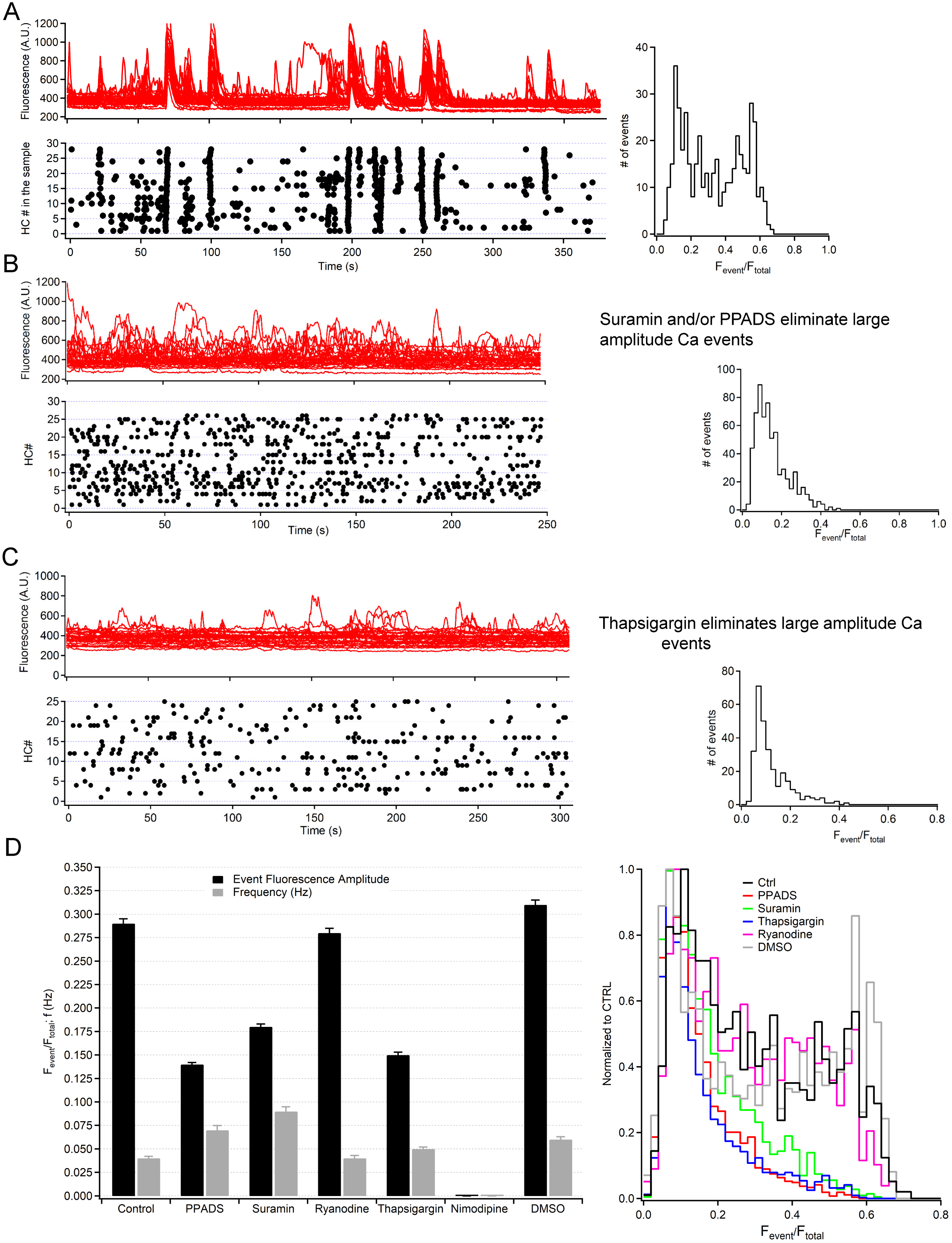
Pharmacological dissection of low- and large-amplitude Ca^2+^ transients in IHCs of mice expressing Ca^2+^ biosensors. **(A)** Representative control records and corresponding amplitude distribution histogram; red traces: raw fluorescence data from a group of contiguous IHCs; black dots: events detected using the algorithm illustrated in **Figure 8. (B)** Effect of suramin. (**C**) Effect of thapsigargin. (**D**) Pooled data quantifying the effects of PPADS (100 µM), suramin (200 µM), ryanodine (50 µM), thapsigargin (10 µM) and nimodipine (50 µM); Ctrl, no treatment; DMSO, control to show lack of effect of the solvent (<0.1%) used for some of the water insoluble tested compounds; Data presented as means ± standard error of means (S.E.M); numbers of samples for each experimental condition are as follows: CTRL, *n* = 6; PPADS *n* = 4; Suramin, *n* = 3; ryanodine, *n* = 3; thapsigargin, *n* = 6; DMSO, *n* = 3. P-values (ANOVA) for each experimental condition relative to control for amplitude of detected events are as follows: PPADS, *P* = 1.34×10^−4^; suramin, *P* = 0.0032; ryanodine, *P* = 0.5549; thapsigargin, *P* =4.7607×10^−5^; DMSO, *P* = 0.8321. P-values (ANOVA) for each experimental condition relative to control for frequency of detected events are as follows: PPADS, *P* = 0.017; suramin, *P* = 0.0016; ryanodine, *P* = 0.7601; thapsigargin, *P* = 0.847; DMSO, *P* = 0.1374.

### Statistical Analysis

Mean values are quoted ± S.E.M. (standard error of the mean). Two or more groups were compared by using two-tailed analysis of variance (ANOVA) or paired *t* test. The Mann-Whitney U test was used for data which were not normally distributed and/or had dissimilar variance. P-values are indicated by letter *P*, where *P* < 0.05 was selected as the criterion for statistical significance. Normality of distribution was determined using the Kolmogorov–Smirnov test.

No statistical methods were used to predetermine sample size. The experiments were not randomized and the investigators were not blinded to allocation during experiments and outcome assessment. No pre-established criteria were used to include/exclude samples or animals.

## RESULTS

### Optical detection of Ca^2+^-dependent APs in mouse cochlear IHCs

Spontaneous Ca^2+^-dependent activity in IHCs and NSCs in the GER was recorded from the apical coil of the immature mouse cochlea (P4-P5) bathed in a perilymph-like extracellular solution (1.3 mM Ca^2+^ and 5.8 mM K^+^: Wangemann, 1996) and at near body temperature. Under the above recording conditions, immature IHCs loaded with the Ca^2+^ indicator Fluo-4 AM exhibited spontaneous and rapid Ca^2+^ transients near their basal (synaptic) region (**Figure 1A,B** and Supplementary Video 1; similar Ca^2+^ transients were present in 10 separate recordings from 9 mice). By combining Ca^2+^ imaging and cell-attached patch clamp recordings, we confirmed that the observed Ca^2+^ signals represent the optical readout of AP activity in IHCs (**Figure 1C**). While individual APs could be detected by Ca^2+^ imaging (**Figure 1C**: left), bursts of APs resulted in an overall increase in the fluorescence level (**Figure 1C**: right) because fluorescence emission increased rapidly, but decayed with an average time constant of 300 ± 11 ms (*n* = 7 APs; **Figure 1D**), which is likely to reflect the dissociation constant (*k*_off_) of the high-affinity Ca^2+^ dye (Canepari and Mammano, 1999; Bortolozzi et al., 2008).

### Intercellular Ca^2+^ waves in the GER propagate along different spatial directions

Using Ca^2+^ imaging we also found spontaneous propagation of intercellular Ca^2+^ waves in the GER, as previously reported (Tritsch et al., 2007; Schutz et al., 2010; Rodriguez et al., 2012; Xu et al., 2017; Mammano and Bortolozzi, 2018; Ceriani et al., 2019). Ca^2+^ waves occurred with a frequency of 3.46 ± 0.44/min in a field of view of 190 µm × 190 µm (10 recordings, 8 cochleae, 8 mice), and can be broadly classified as either *radial* or *longitudinal* (**Figure 2** and Supplementary Video 2). The latter, which correspond to 14% of all waves (**Figure 3A,B,C**), propagated along the coiling axis of the cochlea across the NSCs that intercalate between IHCs with a speed of 7.6 ± 1.7 μm/s (*n* = 7 waves, 4 cochleae, 4 mice). Among the radial Ca^2+^ waves, i.e. those travelling in the direction orthogonal to the coiling axis (86%: **Figure 3B,D,E**), the majority (64%) travelled towards the bulk of the GER (5.7 ± 1.2 μm/s, *n* = 32 waves, 4 cochleae, 4 mice; **Figure 3B,D**) whereas only 22% spread towards the IHCs (5.3 ± 1.2 μm/s, *n* = 11 waves, 4 cochleae, 4 mice; **Figure 3B,E**).

### AP activity in IHCs and Ca^2+^ wave propagation in the GER exhibit variable degrees of correlation

Ca^2+^ imaging revealed that IHCs fired Ca^2+^ APs (as reported optically) also in the complete absence of detectable Ca^2+^ activity in the GER (**Figure 4A,B**). Without Ca^2+^ waves in the GER (e.g. during time interval T1 in **Figure 4B,C**), the IHC-IHC average correlation coefficient (correlation index, *c*, see Materials and Methods) was not significantly different from 0 (*c* = 0.02, *n* = 10 recordings from 9 mice, *P* = 0.1, Student’s t-test, **Figure 4D**), indicating that IHCs generated APs independently of each other. IHC Ca^2+^ activity increased when Ca^2+^ waves were present in the GER (e.g. during time interval T2 in **Figure 4B,C**; see also Supplementary Video 3), leading to an increased probability of synchronized bursting among many hair cells. In these conditions, correlation index values deviated significantly from 0 for 68 out of the 111 Ca^2+^ wave events analyzed (10 recordings from 9 mice, P<0.05, Student’s t-test) and displayed a monotonic dependence with the area of the tissue invaded by the Ca^2+^ wave (Pearson’s coefficient *ρ* = 0.6, **Figure 4D**).

To confirm that AP activity in IHCs and Ca^2+^ wave propagation in the GER can occur independently of each other, we performed simultaneous recording of Ca^2+^ signals from GER NSCs and cell-attached recordings of APs from IHCs (**Figure 5** and Supplementary Video 4). When bursts of IHC AP activity were correlated to Ca^2+^ transients in nearby NSCs (**Figure 5A,B**), their frequency increased significantly (2.8 ± 0.1 Hz, *n* = 82 bursts in 7 IHCs) compared to the condition when no correlation was observed (1.4 ± 0.1 Hz, *n* = 51 bursts in 7 IHCs, *P* < 0.001; **Figure 5C**). AP burst duration was also significantly higher when correlated to non-sensory cell activity (correlated: 11.0 ± 1.5 s; uncorrelated: 2.9 ± 0.3 s, *P* < 0.001, **Figure 5D**). In these conditions, the longitudinal and radial dimensions of the GER area showing significant correlation with AP activity of a given IHC were 310 ± 49 μm and 121 ± 12 μm, respectively (*n* = 7 cochleae; **Figure 6**). This quantitative analysis suggests that GER Ca^2+^ waves promote synchronization of IHC AP activity by variable amounts, depending on wave extension.

Consistent with the observation that a fraction of the Ca^2+^ waves appeared to depart from the IHC region and travelling towards the bulk of the GER (**Figure 4** and Supplementary Video 5), we found that IHCs depolarization triggered Ca^2+^ signals in nearby NSCs, with a delay that was proportional to the distance from the stimulated IHC (3 recordings from 3, **Figure 7**; Supplementary Video 6). Therefore, there appears to be a reciprocal relationship between Ca^2+^ signals generated by IHCs and NSCs of the GER.

### Transgenic mice expressing genetically encoded Ca^2+^ biosensors illuminate AP activity

To gain further insight into the mechanisms that govern spontaneous Ca^2+^ signaling in the developing cochlea, we used transgenic mice (see **Materials and Methods**) that expressed Ca^2+^ biosensors either in the whole sensory epithelium (GCaMP3;Emx2-Cre mice, **Figure 8*A*** and Supplementary Video 7) or selectively in hair cells (GCaMP3;Gfi1-Cre mice, **Figure 8B** and Supplementary Video 8, and GCaMP6f;Gfi1-Cre mice, **Figure 8C** and Supplementary Video 9). The higher contrast and signal-to-noise ratio afforded by cultures obtained from these mice, permitted us to partition IHC Ca^2+^ signals in two broad classes, namely low-amplitude (e.g. those corresponding to downward arrowheads in **Figure 8D**) and large-amplitude Ca^2+^ transients (e.g. downward arrows in **Figure 8D**). Whereas the IHC low-amplitude events were largely uncorrelated, the IHC large-amplitude events occurred only in the presence of Ca^2+^ waves in the GER and were correlated within a group of contiguous IHCs (**Figure 9A**). The two types of signals could be also discriminated pharmacologically. In prior work (Rodriguez et al., 2012) spontaneous Ca^2+^ signaling and Ca^2+^ wave propagation in the GER were reversibly reduced by the phospholipase C (PLC) inhibitor U73122 as well as by the inositol 1,4,5-trisphosphate receptor (IP_3_R) antagonist 2-aminoethoxydiphenyl borate (2-APB), and irreversibly suppressed by thapsigargin, a noncompetitive inhibitor of the Ca^2+^-ATPase (SERCA) that causes the complete and irreversible depletion of Ca^2+^ from the endoplasmic reticulum (ER) Ca^2+^ store. In this work, both Ca^2+^ waves in the GER and IHC synchronous large-amplitude Ca^2+^ transients were abrogated by P2Y and P2X receptor antagonists suramin (200 µM) and PPADS (100 µM), as well as by thapsigargin (10 µM), whereas ryanodine (50 µM) was not effective. However these drugs failed to suppress the desynchronized low-amplitude Ca^2+^ transients in IHCs (**Figure 9B,C**). In contrast, the latter were obliterated by nimodipine (50 µM), whereas Ca^2+^ waves in the GER were able to propagate in the presence of this L-type Ca^2+^ channels inhibitor (Supplementary Video 10). A summary of these results is presented in histogram form in **Figure 9D**. Together, the results in **Figures 8** and **9** confirm that Ca^2+^ waves in the GER require the PIP_2_/PLC/IP_3_R signaling axis promoting Ca^2+^ release from ryanodine-insensitive intracellular Ca^2+^ stores (Rodriguez et al., 2012) and that synchronization of IHC activity by Ca^2+^ waves in the GER depends on extracellular ATP acting on IHCs (Johnson et al., 2011). They also support the tenet that IHC AP firing can occur independently of Ca^2+^ waves in the GER.

### Patch clamp recordings reveal spontaneous APs in isolated IHCs

To confirm that Ca^2+^ APs in immature IHCs do not require Ca^2+^ waves in the GER, we performed whole-cell patch clamp recordings from mechanically isolated IHCs, which were obtained using a small suction pipette to remove the surrounding NSCs. This procedure did not compromise cell integrity, since these isolated IHCs showed normal outward K^+^ currents (**Figure 10A:** amplitude measured at 0 mV: 2.6 ± 0.2 nA, *n* = 3, P2) and a resting membrane potential (−62.0 ± 0.7 mV, *n* = 3) similar to that recorded from IHCs kept *in situ* (Marcotti et al., 2003a; Marcotti et al., 2003b). Importantly, isolated IHCs exhibited spontaneous APs (**Figure 10B**), indicating that this firing activity is an intrinsic property of immature IHCs that can be modulated by the spontaneous Ca^2+^ waves originating from the GER.

**Figure 10.**
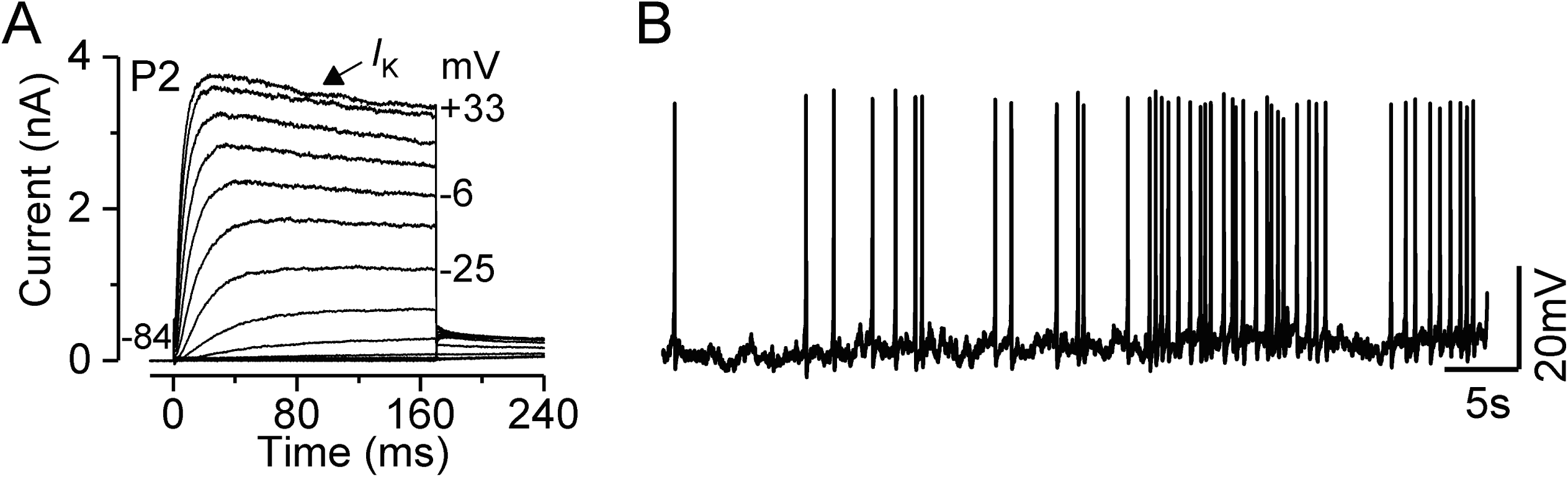
Whole cell recordings from isolated IHCs. **(A)** Typical current responses from an isolated P2 IHCs recorded from the apical coil of the cochlea of wild-type mice. Outward currents were elicited by using depolarizing voltage steps (10 mV increments) from the holding potential of –84 mV to the various test potentials shown by some of the traces. **(B)** Spontaneous action potentials recorded from an isolated P2 apical IHC using whole-cell voltage clamp. IHCs were isolated by using a small pipette to remove the surrounding non-sensory cells.

## DISCUSSION

Using a combination of Ca^2+^ imaging and cell-attached patch clamp recordings performed at body temperature, we have found that AP activity in mouse pre-hearing IHCs is intrinsically generated by the cells themselves. A stringent cross-correlation analysis has revealed that only the Ca^2+^ waves in the NSCs originating within specific distances from IHCs (∼300 µm along the coiling axis of the cochlea and ∼120 µm in the orthogonal, radial, direction) act as an extrinsic pathway to synchronize the intrinsic firing activity of IHCs. We also found that a large fraction of these Ca^2+^ waves originating in the proximity of the IHCs were able to propagate away from the sensory cells towards the bulk of the GER. IHCs depolarization also triggered Ca^2+^ signals in NSCs, which propagated towards the GER, indicating the presence of a bi-directional signaling within the immature organ of Corti. We also found that synchronization of IHC activity by Ca^2+^ waves in the GER depends on extracellular ATP directly acting on IHCs (Johnson et al., 2011). We argue that a combination of intrinsic activity and mutual influence between IHCs and supporting cells forms an intricate feedback mechanism to control the level of AP synchronization in IHCs and generate the patterned activity implicated in the refinement of the auditory pathway before the onset of hearing.

### Nature of spontaneous activity in cochlear non-sensory cells

In the GER, spontaneous Ca^2+^ transients arise periodically, and propagate as Ca^2+^ waves as shown in **Figures 2 to 5**. Our results (**Figures 8** and **9**) support the notion that Ca^2+^ waves in the GER require functional ryanodine-insensitive intracellular Ca^2+^ stores, which are activated by extracellular ATP. Specifically, the binding of ATP to P_2_Y promotes the generation of diacylglycerol and IP_3_ from PLC-dependent hydrolysis of PIP_2_. Intracellular diffusion of IP_3_ and subsequent binding to IP_3_ receptors triggers Ca^2+^ release from the ER, increasing the cytosolic Ca^2+^ concentration to peak levels of ∼500 nM (Beltramello et al., 2005; Ceriani et al., 2016a). This elevated Ca^2+^ concentration has been shown to increase the open probability of connexin hemichannels expressed in NSCs (De Vuyst et al., 2006; Zhang et al., 2006; De Vuyst et al., 2009; Leybaert et al., 2017; Carrer et al., 2018; Hu et al., 2018), which favors the release of ATP from cytosol to extracellular milieu through hemichannels composed of Cx26 and Cx30 protomers (Anselmi et al., 2008; Majumder et al., 2010; Xu et al., 2017; Mammano, 2019). Purinergic signaling is terminated by ectonucleotidases (Housley et al., 2002; Vlajkovic et al., 2004) which degrade extracellular ATP (Zimmermann et al., 2012). Thus, ATP–dependent ATP–release enables the regenerative cell–to–cell propagation of Ca^2+^ signals across the network of cochlear non– sensory cells by exploiting a cascade of biochemical reactions governed by critical phenomena that control also the frequency of cytosolic Ca^2+^ oscillation (Ceriani et al., 2016a).

### Nature of spontaneous activity in cochlear hair cells

Although cochlear hair cells are already visible at E14.5, and exhibit biophysical properties compatible with immature differentiating IHCs (Marcotti et al., 2003a), their maturation into fully functional sensory receptors only occurs at the onset of hearing (Kros et al., 1998).

During pre-hearing stages of development, both IHCs and OHCs are capable of eliciting Ca^2+^-dependent APs (IHCs: (Kros et al., 1998; Beutner and Moser, 2001; Marcotti et al., 2003b; Johnson et al., 2011); OHCs: (Ceriani et al., 2019). Calcium AP activity in apical coil IHCs occurs in bursts-like modality (Johnson et al., 2011; Eckrich et al., 2018). The present results, and a vast body of prior work, indicate that these APs are intrinsically generated by immature hair cells, since they are present in the absence of spontaneous Ca^2+^ waves in NSCs (IHCs, Johnson et al., 2017; Eckrich et al., 2018; Harrus et al., 2018; OHCs, Ceriani et al., 2019), and can be recorded from isolated IHCs (**Figure 10**). This spontaneous firing activity is possible because cochlear hair cells express L-type Ca_v_1.3 voltage-gated Ca^2+^ channels (Platzer et al., 2000; Michna et al., 2003) that activate at about −70 mV (Zampini et al., 2010), which is more negative than their average resting membrane potential (about −65 mV) (Marcotti et al., 2003b). This intrinsic activity has been shown to be modulated by the cholinergic inhibitory medial olivocochlear fibers (Glowatzki and Fuchs, 2000), which form direct, but transient, synapses with immature IHCs (Simmons et al., 1996a; Simmons et al., 1996b). Cholinergic Ca^2+^ signals are controlled and compartmentalized by high levels of intracellular Ca^2+^ buffering and “subsynaptic” cisterns (Moglie et al., 2018). The activation of the efferent system causes immature IHCs to hyperpolarize, thereby reducing the frequency of spontaneous APs (Glowatzki and Fuchs, 2000; Marcotti et al., 2004; Johnson et al., 2011). In mice lacking a functional efferent innervation, both the maturation of the hair cell’s ribbon synapses (Johnson et al., 2013) and the refining of the immature tonotopic maps in the auditory brainstem nuclei (Clause et al., 2014) is impaired. IHC firing activity can also be modulated in the developing cochlea by the mechanoelectrical transducer channel, which is crucial for the maturation of the IHCs (Corns et al., 2018), and by ATP-induced intercellular Ca^2+^ waves from connexin extrajunctional hemichannels in the NSCs in the GER (Tritsch et al., 2007; Schutz et al., 2010; Rodriguez et al., 2012).

Despite the large variability in the direction of propagation, all Ca^2+^ waves reaching the IHC area produced an increased firing activity (long bursts: **Figures 1,5;** see also Eckrich et al., 2018), leading to synchronized firing between nearby IHCs (**Figure 4**). Several data indicate that ATP release from connexin hemichannels in NSCs, which express mainly Cx26 and Cx30, is able to affect IHC activity directly (Tritsch et al., 2007; Johnson et al., 2011), which is also supported by recent observations showing that the application of ATP is able to increase IHC firing prior to the induction of Ca^2+^ waves in the GER NSCs (Eckrich et al., 2018). An alternative mechanism is that ATP, by acting on purinergic auto-receptors expressed in the supporting cells surrounding the IHCs, leads to the opening of TMEM16A Ca^2+^-activated Cl^-^ channels and the efflux of K^+^ and the depolarization of the sensory cells (Wang et al., 2015). However, a recent work has shown that synchronization of the activity of nearby IHCs was also observed in the absence of Ca^2+^ waves in cells of the GER (Eckrich et al., 2018), and suggested instead that it is the efflux of K^+^ from the IHCs during repetitive APs that depolarize neighbouring IHCs and thus coordinating their electrical activity. As mentioned above, we found that prolonged IHC depolarization, which is likely to results in elevation of [K^+^] in the confined extracellular space between IHCs and inner supporting cells (ISCs), triggered Ca^2+^ transients in NSCs (**Figure 7**).

### Further implications for the observed Ca^2+^ signaling activity

Ca^2+^ waves are known to be accompanied by crenation of NSCs, an osmotic effect which determines the transient rupture of cell-cell contacts within Kölliker’s organ following the opening TMEM16A Ca^2+^-activated chloride channels expressed by these cells (Tritsch et al., 2007; Wang et al., 2015). These cellular process correlate with expression of apoptosis and autophagy markers in NSCs of the GER (Hou et al., 2019). Programmed cell death is crucial for animal development and Ca^2+^ signaling contributes to organogenesis, topological organization and orientation (Brodskiy et al., 2019; Fontana et al., 2019) by the interplay of apoptosis and autophagy (La Rovere et al., 2016; Giorgi et al., 2018). Therefore, we speculate that, besides modulating AP firing in IHCs, Ca^2+^ waves are likely to support the tightly regulated autophagic and apoptotic processes required for the development and formation of the mature cochlear sensory epithelium. Indirect support for this hypothesis is offered, for example, by the evidence that normal hair cell development is impaired in knockout mice for Cx26 and Cx30 (Johnson et al., 2017), which is followed by the degeneration of both hair cells and NSCs (Crispino et al., 2011). These Cx deficient mice were also shown to have increased levels of oxidative stress in the cochlea due to impairment of Nrf2/ARE signaling, accelerated age-related hearing loss (Fetoni et al., 2018). In multiple cell types, oxidative stress potentiates IP_3_–receptor 1 (IP_3_R1)–mediated Ca^2+^ release by oxidation of numerous cysteine residues (Joseph et al., 2018), and disturbed Ca^2+^ signaling has the potential to derail the course of autophagy and cell death (La Rovere et al., 2016).

Summarizing, it has been very effectively pointed out that “each Ca^2+^ wave is a specific and unique code that is indispensable for the transfer of important information based on its precise amplitude, frequency, timing, and duration” (Giorgi et al., 2018). The Ca^2+^ signaling described in this article sits at a crucial crossroad between development and its derailment, autophagy, apoptosis and premature aging of the mammalian auditory system.

## Acknowledgements

This work was supported by Fondazione Telethon (grant GGP13114 to FM), the University of Padova (Progetto Strategico DYCENDI, prot. STPD11ALFE, to FM), The Wellcome Trust (grant 102892 to WM) and NIH intramural research funds (Z01-DC000002 to B.K.). FC was partially supported by a junior post-doctoral fellowship from the University of Padova (grant CPDR132235 to FM). SLJ is a Royal Society URF.

